# Differential stimulation of the retina with glutamate toward a neurotransmitter-based retinal prosthesis

**DOI:** 10.1101/060681

**Authors:** Corey M. Rountree, Samsoon Inayat, John B. Troy, Laxman Saggere

## Abstract

Subretinal stimulation of the retina with neurotransmitters, the normal means of conveying visual information, is a potentially better alternative to electrical stimulation widely used in current retinal prostheses for treating blindness from photoreceptor degenerative diseases. Yet, no retinal stimulation study exploiting the inner retinal pathways exists. Here, we demonstrate the feasibility of differentially stimulating retinal ganglion cells (RGCs) through the inner nuclear layer of the retina with glutamate, a primary neurotransmitter chemical, in a biomimetic way. We show that controlled pulsatile delivery of glutamate into the subsurface of explanted wild-type rat retinas elicits highly localized simultaneous inhibitory and excitatory spike rate responses in OFF and ON RGCs. We also present the spatiotemporal characteristics of RGC responses to subretinally injected glutamate and the therapeutic stimulation parameters. Our findings could pave the way for future development of a neurotransmitter-based subretinal prosthesis offering more naturalistic vision and better visual acuity than electrical prostheses.

## Introduction

Photoreceptor (PR) degenerative diseases such as retinitis pigmentosa and macular degeneration are leading causes of presently incurable blindness. The irreversible vision loss in these diseases is ultimately due to the loss of PR cells, which normally transduce visual information into chemical signals that activate the retinal ganglion cell (RGC) layer through a network of neurons in the inner nuclear layer (INL). Of the various strategies explored for treatment of PR degeneration over the last two decades, including gene therapy and stem cell transplantation, retinal-based prostheses have emerged as the most promising to restore vision in blind patients^1^. The treatment strategy of retinal prostheses is predicated on artificial stimulation of either RGCs or INL neurons. Direct stimulation of RGCs with a prosthesis in the epiretinal (front of the retina) configuration bypasses INL neurons, while INL stimulation with a prosthesis in the subretinal (behind the retina) configuration seeks to exploit the inherent visual processing capability of INL neurons, which remain intact and functional despite the significant retinal remodeling subsequent to PR degeneration^1,2^. The latter scheme offers the possibility of differentially stimulating the OFF and ON pathways of the retina at the synaptic layer where the separation between these parallel pathways originates. These parallel pathways separate visual stimuli into OFF and ON components (e.g. a point of light would excite the ON pathway and inhibit the OFF) and play a major role in processing and mediating contrast sensitivity in visual perception^3^. Since the distinction between the OFF and ON components is maintained throughout the entire visual system, differential stimulation of these pathways is critical to restoring naturalistic vision to patients with PR degeneration with a retinal prosthesis. In addition, a retinal prosthesis must be able to localize the stimulation ideally to a single cell or a few cells to achieve high spatial specificity, i.e., good visual acuity.

Presently, all existing subretinal prostheses designed for stimulating INL neurons employ electrical current to stimulate target neurons. Though electrical stimulation has been optimized to evoke functional RGC responses, electrical current activates all types of retinal cells including their axons indiscriminately^4,5^. This non-selective nature of electrical stimulation results in the simultaneous excitation of both the OFF and ON pathways in the INL, creating unnatural and confusing perceptions in patients^5,6^. Although some groups are actively exploring techniques to generate quasi differential stimulation effects in the retina with electrical current^7,8^, electrical-based prostheses are unlikely to restore naturalistic vision since this form of quasi differential stimulation does not engage INL circuitry. Another major drawback of electrical stimulation is that it requires large-diameter electrodes to safely handle the high currents required to stimulate degenerated retinas without tissue damage^1,5^. This has limited the visual acuity that electrical prostheses in clinical use today can restore to levels (LogMARs 1.43-2.9) much worse than the legal definition of blindness (LogMAR 1.0)^9^. Thus, at present, restoration of high resolution naturalistic vision with electrical-based prostheses remains elusive despite significant advancements in retinal prosthesis technology over the past two decades^9–12^, and therefore, it is worthwhile to explore alternative agents that could stimulate the retina more effectively along established visual pathways.

A neurotransmitter chemical, particularly glutamate, the main agent of intercellular communication in the normal retina, has long been suggested as a more effective stimulus agent than electrical current for a retinal prosthesis, though very few studies have explored the feasibility of retinal stimulation with neurotransmitters^13–16^. Two studies^15,16^ that have demonstrated the feasibility of glutamate stimulation of in-vitro preparations of whole retinas focused on direct stimulation of RGCs from the epiretinal side. These studies showed glutamate to be effective and advantageous over electrical current in stimulating RGCs, but since they bypassed INL neurons, they did not take advantage of the intrinsic visual processing capability of INL circuitry. In the normal retina, visual information processed by photoreceptors is separated and transmitted to the OFF and ON pathways at photoreceptor-bipolar cell synapses through the release of glutamate onto a variety of glutamate receptors in the outer plexiform layer (OPL)^17^. Evidence from numerous studies on retinal degeneration modeling indicates that INL cell types continue to exhibit glutamatergic responses even in late stages of degeneration, although the glutamate sensitivity may be reduced or eliminated in some cases^2,18–23^. Theorizing that the loss of glutamate sensitivity is likely caused by the lack of glutamatergic input from PRs, these studies suggest that early therapeutic intervention of a degenerated retina could preserve glutamate sensitivity in INL neurons whereas treating late stage degeneration may be possible, albeit difficult. Based on this evidence, it has been hypothesized that subretinal glutamate stimulation of the INL of a degenerated retina could activate RGCs through the amacrine-bipolar-RGC pathway and allow for differential stimulation of the OFF and ON pathways of visual circuitry^24^. The possibility of activating RGCs through INL neurons has a paradigm-shifting implication for retinal prosthesis technology because this biomimetic stimulation of the retina could restore high resolution naturalistic vision to patients with PR degeneration. Yet, no experimental study on subretinal chemical stimulation has been reported.

Here we address the feasibility of differentially stimulating the retina with glutamate from the subretinal side and present the spatiotemporal characteristics of the glutamate-evoked RGC responses obtained by setting up an experimental platform that allows easy access to both the front and back sides of explanted retinas. Using this platform, we convectively injected controlled volumes of 1.0 mM glutamate 70 μm into the subsurface of in-vitro preparations of wild-type rat retinas via 10 μm-diameter tip micropipettes while recording the RGC responses on the other side with a multielectrode array (MEA). Glutamate was focally delivered over each RGC receptive field (RF) center using a pulsatile pressure of 0.69 kPa and 10-30 ms pulses. The RGC RF centers were determined a priori by stimulating the retinas with Gaussian white noise (GWN) checkerboard stimuli and conducting spike-triggered averaging (STA) of the recorded spike trains.

## Results

### Differential stimulation of RGCs through INL neurons

To investigate if subretinally injected glutamate stimulated RGCs through INL neurons, we analyzed multiple sets of MEA-recorded data comprising RGC spikes evoked by GWN and glutamate stimuli from 18 retina samples. First, we identified a total of 1654 spiking units from all retina samples through spike sorting and then classified populations of RGC subtypes viz., OFF, ON, ON-OFF, or visually nonresponsive, on the basis of their spike responses to GWN stimulation. Next, we examined RGC spike responses to glutamate stimulation from all retinas and found that 551 (of the total 1654) units were responsive to glutamate. The glutamate-responsive units were found to be present across all RGC subtypes, including cells that did not exhibit robust visual responses to GWN (Table 1a), which indicates that glutamate likely stimulated some RGC subtypes, such as directionally selective cells, that could not be easily identified through GWN stimulation^25^. An analysis of the peristimulus time histograms (PSTHs) of glutamate-responsive units revealed that glutamate evoked both excitatory and inhibitory responses in RGCs (see Supplementary Fig. S1) with some RGC subtypes showing a preference toward one or the other (Table 1b). ON RGCs exhibited almost exclusively excitatory responses, while OFF RGCs displayed a mix of purely excitatory, purely inhibitory or both excitatory and inhibitory glutamate responses. Furthermore, OFF RGCs switched their response behavior (i.e., from excitatory to inhibitory or vice versa) as the glutamate injection site moved relative to their RF centers (Fig. 1), with significantly (*p* < 0.01, two-proportion z-test) higher likelihood of cells expressing inhibitory responses to injections close to their RF centers and excitatory responses to injections farther away.

The presence of inhibitory responses to glutamate, primarily from OFF RGCs, demonstrates indirect stimulation of RGCs through other presynaptic neurons because RGCs have strictly excitatory responses to glutamate^17^. Excitatory responses, however, might result from either direct activation of RGC synapses and/or those of presynaptic neurons. Furthermore, we observed differential responses (i.e., simultaneous excitatory ON and inhibitory OFF RGC responses to a given stimulus) in about 60% (66 out of 114 sets) of the glutamate injections that elicited both OFF and ON RGC responses (see Fig. 2) and purely excitatory responses suggestive of direct activation of RGCs in the remaining 40% of glutamate injections. These observations, specifically the high proportion of simultaneous differential responses, strongly demonstrate that exogenous glutamate can differentially stimulate the OFF and ON subsystems and imply that glutamate can modulate RGC firing through different retinal neural pathways, most likely INL neurons.

**Table 1.** Populations of RGC sub-types responsive to subretinal glutamate injections and their response polarity. (**a**) Number of glutamate-responsive (GR) and glutamate-nonresponsive (GNR) RGCs classified into subtypes. GNR units were mostly recorded on electrodes far from the injection site. (**b**) Classification of GR RGCs based on their response polarity (excitation, inhibition, or a mix of both, i.e., excitation to one set of injections and inhibition to another). ON RGCs showed a strong preference towards excitatory responses while roughly half of all OFF RGCs displayed some inhibition. ON-OFF and unclassified cells exhibited a strong preference toward excitation though also presented inhibitory or mixed responses. When mixed, the change in polarity creates a center-surround receptive field, with inhibitory responses evoked centrally and excitatory responses evoked peripherally.

| **a.** | **RGC subtypes** |  |  |
| --- | --- | --- | --- |
|  | **GR** | **GNR** | **Total** |
| **ON** | 60 | 53 | 113 |
| **OFF** | 86 | 101 | 187 |
| **ON-OFF** | 221 | 458 | 679 |
| **Unclassified** | 184 | 491 | 675 |
| **Total** | 551 | 1103 | 1654 |

**Table 1.** Populations of RGC sub-types responsive to subretinal glutamate injections and their response polarity.
| **b.** | **Response polarity** |  |  |
| --- | --- | --- | --- |
|  | **Excitation** | **Inhibition** | **Both** |
| **ON** | 55 (91.7%) | 4 (6.7%) | 1 (1.7%) |
| **OFF** | 42 (48.8%) | 26 (30.2%) | 18 (20.9%) |
| **ON-OFF** | 166 (75.1%) | 30 (13.6%) | 25 (11.3%) |
| **Unclassified** | 125 (67.9%) | 35 (19.0%) | 24 (13.0%) |
| **Total** | 388 (70.4%) | 95 (17.2%) | 68 (12.3%) |

**Figure 1.**
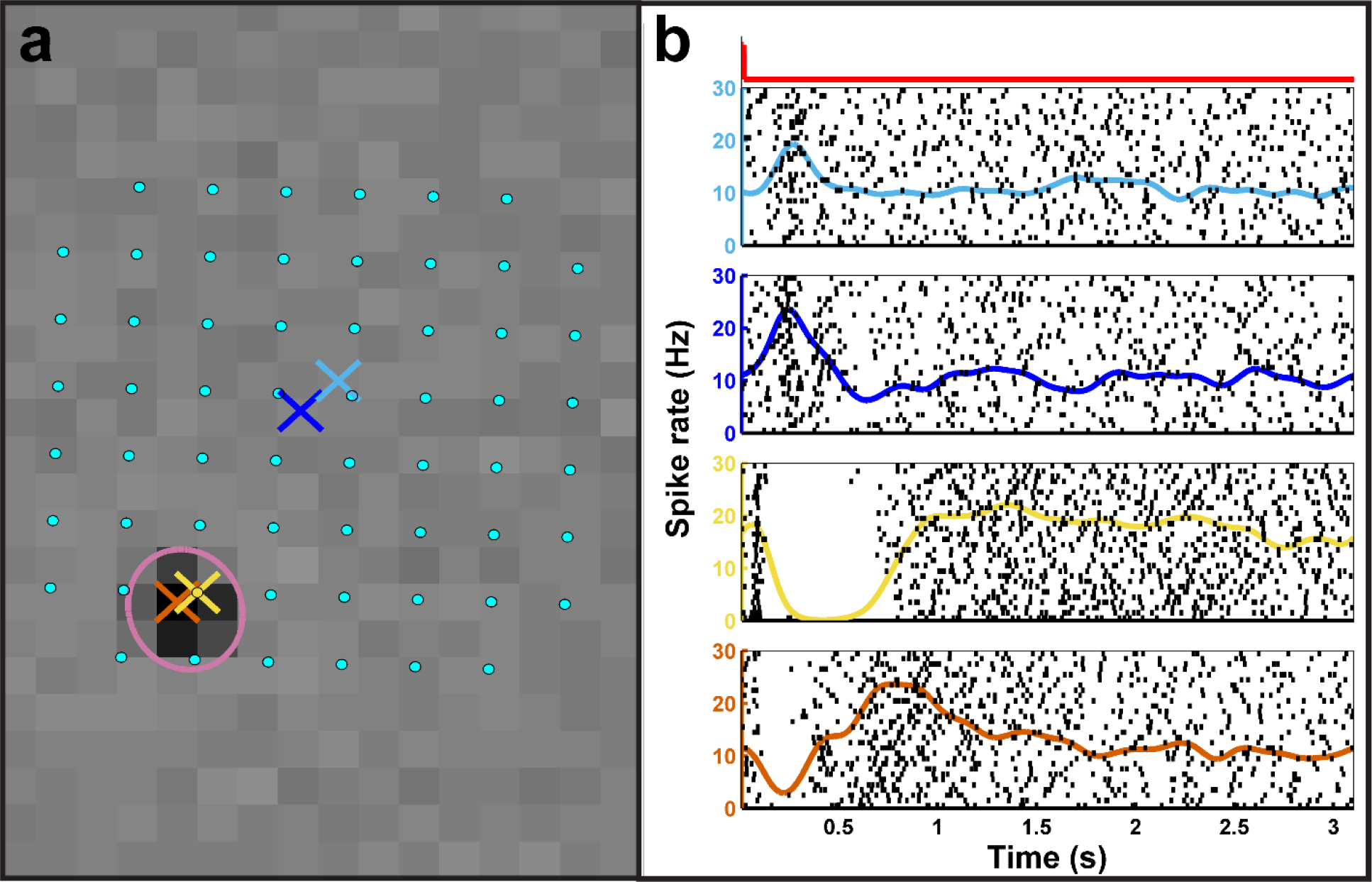
A subset of OFF RGCs responding to glutamate injections displayed both excitatory and inhibitory responses dependent on the distance separating the injection site from their RF centers. (**a**) A map showing the locations of MEA electrodes (cyan dots; 200 p,m separation) and the locations of four glutamate injections (colored ‘×’s) with respect to the RF center of a representative OFF RGC (magenta ellipse) superimposed over this cell’s peak STA frame.(**b**) The spike rate and raster plots showing this cell’s responses to the glutamate injections corresponding to the ‘×’s in the spatial map arranged in descending order from farthest to closest. The injection timing and duration for all glutamate injection events is indicated by the red square wave above the topmost plot on the left. As can be seen, injections far from the RF center elicited excitatory responses while those in or near the RF center evoked inhibition. Similar patterns were seen in approximately 25% of glutamate-responsive OFF RGCs.

**Figure 2.**
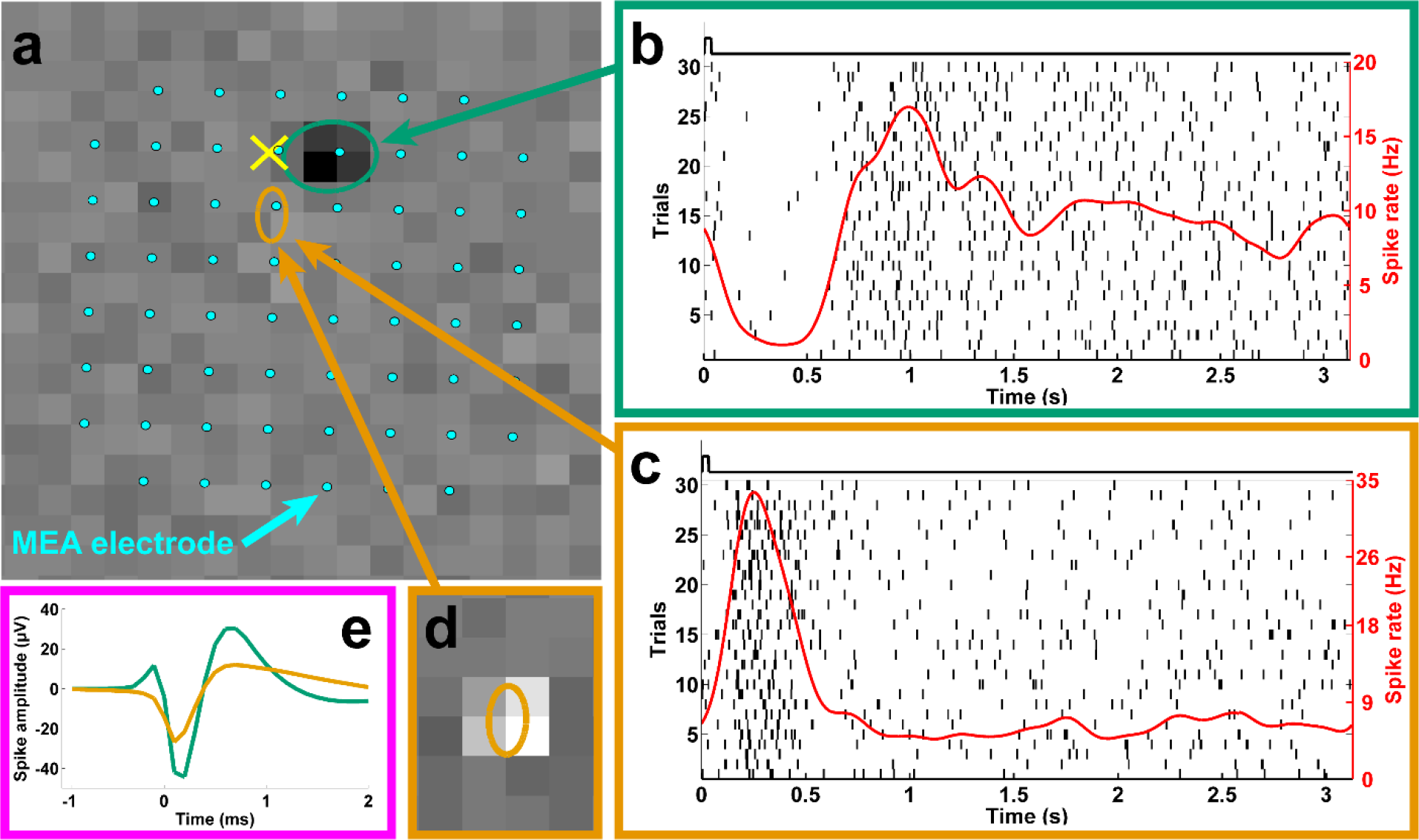
Subretinal glutamate injections elicit colocalized and differential responses in the OFF and ON pathways. (**a**) The locations of the receptive field centers for an OFF (green ellipse) and an ON (orange ellipse) RGC relative to the injection site (yellow ‘×’) overlaid with 60 electrodes of the MEA (cyan dots). The background displays the visually-evoked spike-triggered-average (STA) for the OFF RGC. (**b**) and (c) The raster (vertical black lines) and spike rate (red traces) for each cell. The black square waves above each plot indicate the timing and duration of each glutamate injection event. The ON RGC (**c**) displayed excitation while the OFF RGC (**b**) exhibited an inhibitory response to the same stimulus. (**d**) The STA for the ON RGC. (**e**) Two unique spike shapes corresponding to the two units, which confirms that these are two distinct cells. The different responses of OFF and ON RGCs to a common stimulus strongly suggests differential stimulation of the OFF and ON pathways.

To confirm our inference that subretinal glutamate injections elicited differential responses through INL neural stimulation, we also examined local field potentials (LFPs) recorded with MEA electrodes near injection sites and found large amplitude transients (Fig. 3a). Previous research indicates that LFPs originate from INL neurons^26,27^. The glutamate-evoked LFP (gLFP) transients we observed were triphasic with an initial rising phase (Fig. 3b) and usually preceded (Fig. 3c) significant spike rate responses (gSRs) in nearby units. We recorded LFP data in a subset of recordings and identified 127 sets of glutamate injections that yielded significant transient gSRs, of which 89 (70%) also induced significant gLFPs. The presence of substantial gLFP activity, combined with the high proportion of inhibitory gSRs, supports the conclusion that subretinally injected glutamate modulates INL neurons, such as bipolar or amacrine cells, because RGC activity has been shown to only induce slight amplitude and temporal alterations to light-evoked LFPs, also known as electroretinograms (ERG)^27^.

**Figure 3.**
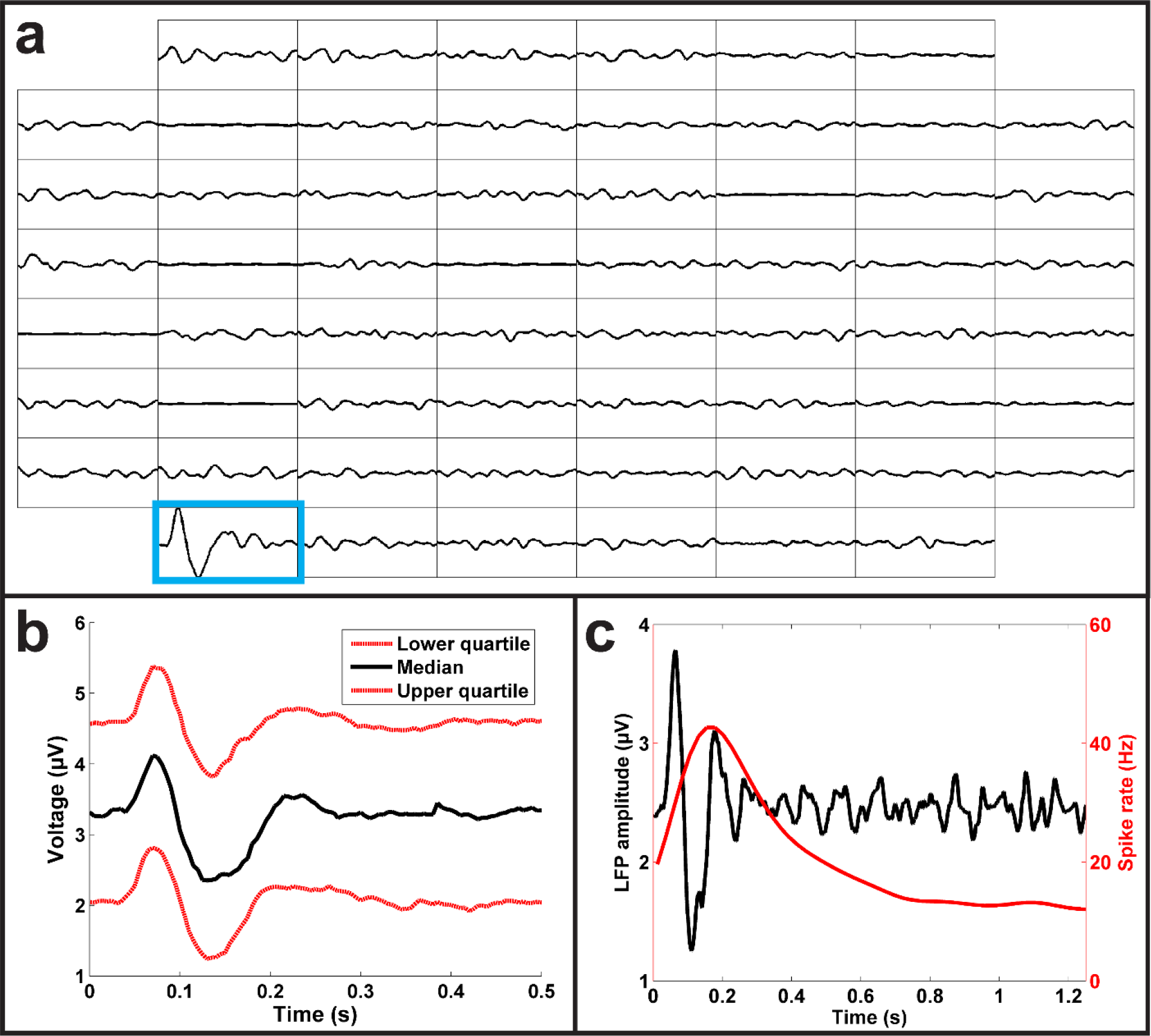
Subretinal glutamate injections elicited robust LFP responses. (**a**) Representation of LFPs recorded from each of the 60 electrodes (interelectrode distance of 200 μm). The only trace that shows a significant LFP response (outlined in blue) corresponds to the electrode where glutamate was injected. (**b**) The lower (bottom red), median (black), and upper (top red) quartile gLFP responses show a characteristic triphasic shape. (**c**) An overlay of the LFP (black) and spike rate (red) responses from a representative ON RGC, demonstrating that LFP responses preceded spike rate responses.

### Spatial resolution of glutamate-evoked responses

As mentioned above, achieving stimulation with high spatial specificity (i.e., high spatial resolution) is vital to restoring good vision with a retinal prosthesis. We define spatial specificity as the spatial extent over which a bolus of glutamate stimulates RGCs, evoking a significant response in either gSRs or gLFPs. We investigated the spatial specificity of subretinal glutamate stimulation by quantifying the distances of RF centers of all glutamate responsive RGCs from the injection site. Fig. 4a shows a color plot of the spatial distribution of gSRs to injections, where the midpoint represents the injection site, warmer colors indicate higher densities of cells responding, and cooler colors indicate regions with fewer responsive cells. The plot reveals the median distance between RF centers and the injection site to be 180 μm and half of all gSRs were confined to distances ranging between 75-310 μm (lower-upper quartiles) from the glutamate injection site. Because the MEA permits only a sparse sampling of RGCs, we fit the data of Fig. 4a with a 2D Gaussian spread function to create a more continuous and presumably accurate estimate of the spatial distribution of stimulation (Fig. 4b). This spread function fits the data well (*r*^2^ = 0.99) and predicts that half of all glutamate-responsive RGCs lie within 230 μm of the injection site. Similarly, we plotted the spatial distribution of gLFP responses (Fig. 4c) and a Gaussian distribution (Fig. 4d) of all gLFP responses to the same glutamate stimulation. The gLFP spatial distribution plot (Fig. 4c) reveals the median distance between electrodes with gLFP responses and the injection site to be 245 μm with half of all gLFPs confined to distances ranging between 110-575 μm (lower-upper quartiles) from the glutamate injection site. The LFP data were also well fit by a Gaussian spread function (Fig. 4d) and revealed the median distance of all glutamate-responsive LFPs to be 105 μm.

**Figure 4.**
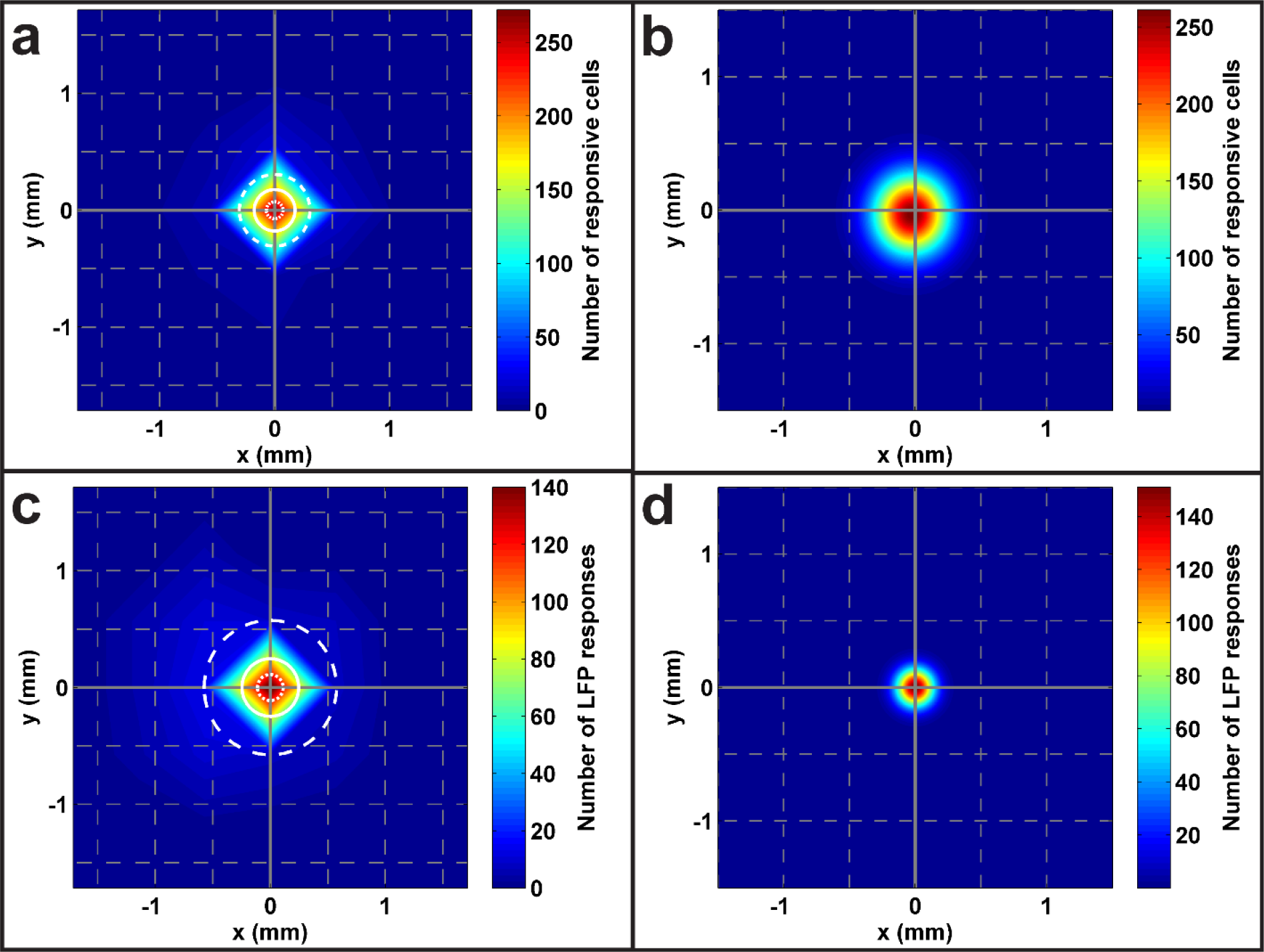
Subretinal injections elicit localized RGC responses. (**a**) A two-dimensional histogram displaying the location of all RGC receptive field centers relative to the injection site (midpoint) that were responsive to glutamate injected below the photoreceptor surface. The solid white circle indicates the median distance of 180 μm while the inner and outer dotted/dashed circles specify the lower (75 μm) and upper (310 μm) quartiles, respectively. Warmer colors represent higher densities of RGCs as quantified by the color bar on the right. The grey gridlines are separated by distances of 500 μm. (**b**) The data in (**a**) were fitted with 2D Gaussian functions to produce a more continuous spatial distribution than predicted in (**a**). The Gaussian fit predicts a median spread of 230 μm for subretinally injected glutamate. (**c**) and (**d**) Two-dimensional histograms showing the spatial distribution and Gaussian fit for LFP responses, which yielded median distances of 245 and 105 μm, respectively.

While characterizing the spatial specificity of glutamate responses, it is important to also consider the response amplitudes as the amplitudes vary with the distance from the injection site. To investigate the efficacy of glutamate stimulation around the injection site, we compared the gSR amplitudes normalized by the peak amplitude of full-field flash-evoked responses for all RGCs in the vicinity of the chemical delivery site. The amplitudes were normalized to allow for population-level comparisons across RGCs because the amplitudes of spike rate responses to glutamate and light stimuli can vary widely from cell to cell. Unsurprisingly, we found that RGCs expressed the largest normalized response amplitudes to injections close to their RF centers (< 200 μm), where they exhibited response amplitudes roughly equivalent to their peak full-field flash amplitudes. Injections farther from the RF center yielded progressively and significantly (*p* << 0.001, Kruskal-Wallis) smaller normalized response amplitudes (Supplementary Fig. S2a). An example of this relationship can be seen in Supplementary Fig. S2b, which shows PSTH and raster plots for a representative RGC in response to injections at several distances from the injection site, indicating the decrease in response amplitude with distance. A similar trend was observed for gLFP amplitudes (Supplementary Fig. S2c). These data suggest that effective RGC responses evoked by 1 mM glutamate injected into the retina at 0.69 kPa for 10-30 ms is spatially localized within a distance of about 200 μm from the injection site. The stimulation effect decreases with increasing distances of RGCs from the injection site, possibly due to decrease in lateral spread of glutamate as well as robust natural mechanisms for glutamate removal in the retina^28^.

### Temporal resolution of glutamate-evoked responses

Any functional retinal prosthesis must be able to provide stimulation that is temporally naturalistic to allow patients to detect motion, especially during activities like locomotion. Here, we define temporal resolution as the highest rate of subretinal glutamate injections that evokes a significant response in either gSRs or gLFPs. We studied gSR responses of 21 RGCs and gLFP responses at three electrodes corresponding to varying injection frequencies from 0.25 Hz to 6 Hz. Fig. 5 shows PSTH plots for a representative RGC (Fig. 5a) and the corresponding LFPs (Fig. 5b) for multiple frequencies of injections. RGCs responded with gSRs to injection frequencies up to 3.5 Hz, but robust gLFP responses corresponding to injection frequencies as high as 6 Hz were recorded. The results suggest that glutamate may modulate the activity of INL neurons at higher frequencies, but cannot drive the RGC firing rates at those higher frequencies. We speculate that this difference between gSR and gLFP response frequencies may be due to the relatively low temperature (22°C) at which this study was conducted because previous work has shown that the latencies of LFP responses are less affected by shifts in temperature than spike rate responses^29^. Extrapolating our data to body temperature (37°C) using the Q10 values reported by Schellart *et al*.^29^, we predict that gSR temporal resolutions as high as 14 Hz may be achievable.

**Figure 5.**
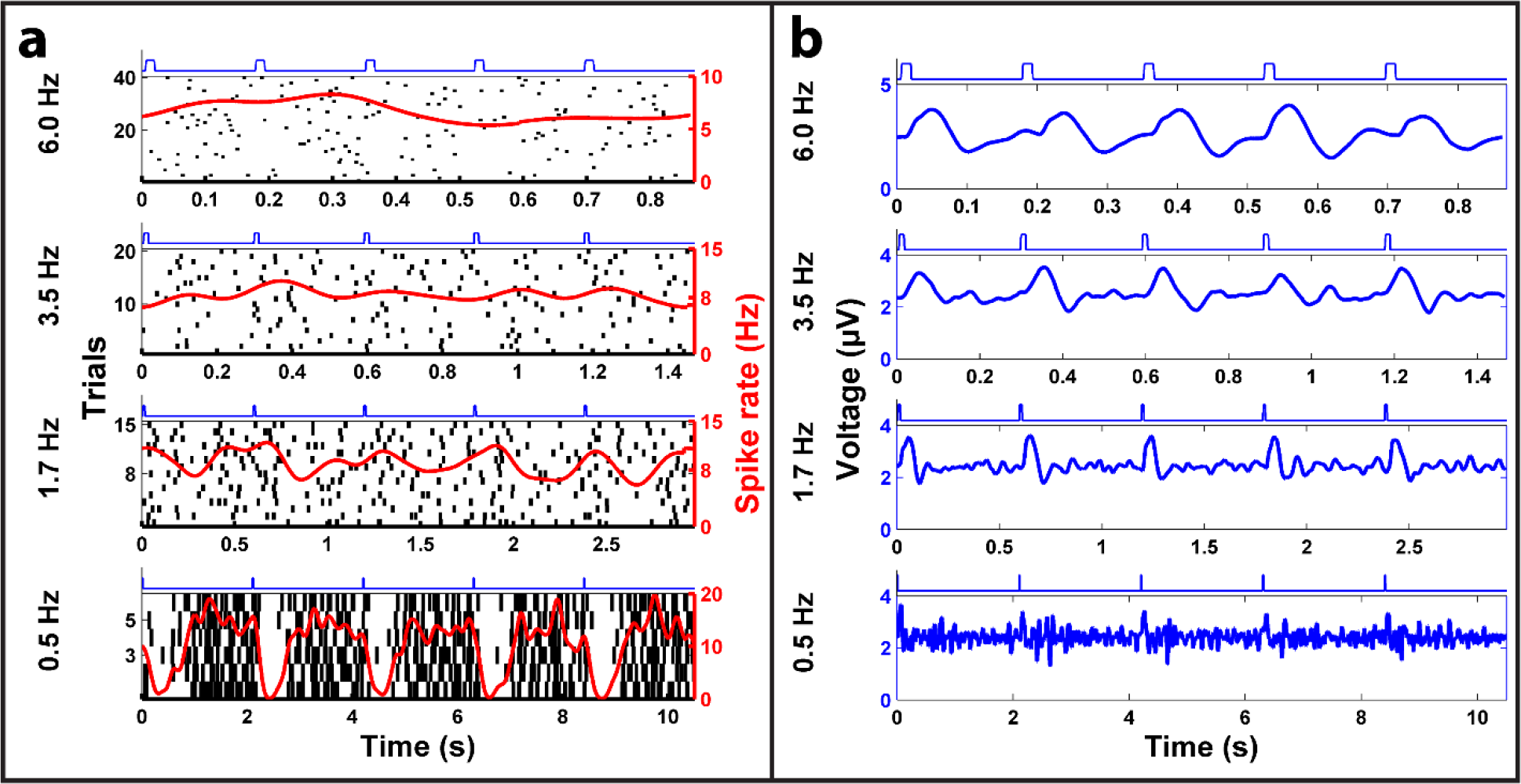
Frequency responses of subretinal glutamate stimulation. (**a**) Representative raster plots of spike discharge for an OFF RGC. (**b**) LFP plots recorded from an electrode near the OFF RGC in response to glutamate injections delivered at four different frequencies (indicated on the far left). Each plot shows the average responses to multiple trials, where each trial consisted of 5 consecutive 10 ms glutamate injections whose timing is represented by the blue square waves above each subplot. Glutamate injections delivered as fast as 3.5 Hz produced discernable spike rate responses, while injections at the highest frequency (6 Hz) tested elicited robust LFP responses.

## Discussion

Pulsatile injections of glutamate, the primary retinal neurotransmitter chemical, delivered into explanted rat retinas at the PR-bipolar cell synapse layer (approximately 70 μm below the PR surface) where the ON and OFF pathways are first established, elicited robust physiological responses from RGCs. After exploring the therapeutic ranges of the key stimulation parameters, viz., the glutamate concentration, injection pressure, pulse width and frequency, we determined an optimal set of the stimulation parameters to accomplish the three most important stimulation traits for a neurotransmitter-based subretinal prosthesis: biomimetic differential stimulation, spatially localized stimulation and temporally naturalistic stimulation. We found that small boluses of 1.0 mM glutamate convectively released into the INL of the retina using 0.69 kPa and 10-30 ms pressure pulses can differentially stimulate retinal OFF and ON pathways, eliciting colocalized inhibitory and excitatory responses in OFF and ON RGCs, respectively.

While exogenous glutamate released in the OPL was theorized to mimic the retina’s natural synaptic communication and differentially activate the ON and OFF pathways, the therapeutic set of parameters necessary for practically achieving differential stimulation were unexpected given the multiplicity of glutamate-responsive neuron types throughout the retina^17^. To examine the pathways by which glutamate differentially stimulated OFF and ON RGCs, we sought to narrow down the cellular targets of the exogenous glutamate in the INL. OFF and ON bipolar cells, which establish the distinction between these pathways in the OPL, were the ideal intended targets to produce differential stimulation, but the presence of almost exclusively excitatory ON responses combined with a mix of excitatory and inhibitory OFF responses suggests that the injected glutamate most likely engaged an unexpected and more complex set of retinal circuits. Based on our results, we can exclude stimulation of cell types that cause inhibitory ON responses such as ON bipolar, photoreceptor, and horizontal cells^17,30,31^. In fact, an examination of the retinal pathways reveals that glutamate should only yield inhibitory OFF responses (without inhibitory ON responses) if it acted on amacrine cells, most likely the AII cell that is particularly dominant in the rodent retina^32,33^. The AII cell bridges the rod and cone pathways by receiving glutamatergic input from rod bipolar cells and conveying this information to ON bipolar cells through sign-conserving electrical gap junctions and OFF bipolar cells via inhibitory (sign-inverting) glycinergic synapses^33^. When excited with glutamate, AII cells would therefore cause excitatory responses in neighboring ON bipolar cells and inhibit nearby OFF bipolar cells, which corresponds well with our differential response data. In the remainder of stimuli where differential responses were not observed, the excitatory OFF responses could be the result of OFF bipolar cell stimulation, but the simultaneous and colocalized excitation of ON RGCs is interesting and suggests either direct RGC stimulation or a mix of OFF bipolar and AII cell stimulation.

Additional work would be required to convincingly identify the cellular targets for differential stimulation and localize glutamate delivery to those target cells. Nevertheless, our results demonstrate that subretinal glutamate injections can reliably elicit differential responses in the OFF and ON pathways through stimulation of INL neurons, with evidence pointing towards AII cell activation.

The spatial specificity (lower and upper quartile distributions of glutamate-responsive RGCs) achieved with single-port glutamate stimulation of rat retinas in this study, if reproduced in human retinas, correspond to visual acuities in the range 1.2-1.8 LogMAR, which compare favorably with those reported for electrical prostheses in clinical use today. We speculate that even better visual acuities closer to the legal definition of blindness may be possible with multi-site glutamate stimulation using a large array of independently addressable injection ports, each providing spatially localized stimulation. Furthermore, the maximum temporal stimulation rate of 3.5 Hz demonstrated at 22°C with pulsatile glutamate injections is comparable to what has been deemed adequate for enabling rudimentary reading and daily functions in patients using electrical prostheses in clinical trials^12^. Higher temporal stimulation rates are likely achievable with finer on-chip actuation of glutamate in a device interfaced with the retina at physiological temperatures^34–36^.

Although an exogenous glutamate stimulation strategy is ultimately targeted for retinas with degenerated PRs, we purposely conducted this study on wild-type rat retinas (with functional PRs) in order to systematically guide the study by identifying RGC subtypes and their RF centers with GWN checkerboard stimulation. Determination of RGC subtypes via GWN checkerboard stimulation facilitated the analysis of gSRs to confirm whether or not differential stimulation was achieved with a particular glutamate injection event. Identification of RGC RF centers enabled fine characterization of the spatial localization of glutamate responses in terms of distances between the glutamate-responsive RGCs and the injection site. Further work would be required to reproduce the findings of this study in degenerated retinas. Nevertheless, since both bipolar and AII cells are known to preserve some level of glutamate sensitivity even in late stages of degeneration^2^, we conjecture that the results and conclusions of this work apply to a degenerated retina, though glutamate stimulation efficacy may be reduced depending on the level of retinal degeneration.

Our findings are encouraging for advancing the development of a neurotransmitter-based subretinal prosthesis as a serious and more effective alternative to electrical-based prostheses for restoring useful vision to patients with PR degenerative diseases. A neurotransmitter-based subretinal prosthesis could potentially not only reduce the burden for external visual signal processing electronics, but also restore more natural vision to patients by taking advantage of the retina’s inherent visual processing circuitry. We envision that a neurotransmitter-based retinal prosthesis could be enabled by fabricating a MEMS (microelectromechanical-systems)-based microfluidic device featuring a high density array of independently addressable hollow microneedles in the subretinal space that store and release pulsatile boluses of neurotransmitters near target INL synapses (see Supplementary Fig. S3). To develop such a device, a number of key parameters, including the device geometry (e.g., port diameters and spatial resolution) and chemical delivery parameters (e.g., range of pressures and pulse widths of glutamate injection), must be optimized. Although the stimulation parameters would have to be optimized for the type and stage of retinal degeneration, the optimal set of glutamate stimulation parameters determined here offers baseline parameters to guide future work towards the development of a neurotransmitter-based retinal prosthesis.

## Methods

### Study Design

The study design was based on controlled laboratory experiments involving in-vitro stimulation of explanted rat retinas with glutamate. The objective of the study was to convectively inject pulsatile boluses of glutamate chemical into the subsurface of retinas to evoke physiological responses comparable to the normal light-evoked responses. Based on the findings of several pilot experiments carried out to investigate therapeutic ranges of various stimulation parameters (glutamate concentration, injection pressure, pulse width and frequency), an optimal set of therapeutic stimulation parameters (1 mM glutamate, 0.69 kPa, 10-30 ms, 0.25-6 Hz) that elicited robust glutamate-evoked retinal responses was employed in this study.

Neurotransmitter-based stimulation was systematically investigated using a sample size of 18 retinas, which was determined by the total number of glutamate-responsive RGCs identified across all stimulation trials in this study. Data collection was completed once a total of at least 500 glutamate-responsive RGCs were identified to provide a sufficiently large representative dataset for meaningful analyses and conclusions. The only data excluded from data analyses were responses from RGCs with abnormal spike shapes (i.e. responses not quantitatively determined to be from either somal or axonal).

All animal experiments were conducted in accordance with the guidelines outlined by the National Research Council’s *Guide for the Care and Use of Laboratory Animals*. Animal handling and euthanasia protocols were reviewed and approved by the Institutional Animal Care and Use Committee of the University of Illinois at Chicago.

### Retinal Sample Preparation

Retinas were explanted from Hooded Long-Evans rats (PND 25-35, *N* = 18, either male or female; Charles River Laboratories, Wilmington, MA) after they were dark-adapted for at least one hour and euthanized by carbon dioxide followed by cervical dislocation. The sample size (18 retinas) was determined by the total number of glutamate-responsive RGCs identified across all stimulation trials in this study. Data collection was completed once a total of at least 500 glutamate-responsive RGCs were identified to provide a sufficiently large representative dataset for meaningful analyses and conclusions. The explanted retinas were placed onto a perforated MEA (pMEA200/30iR-Ti, Multichannel Systems, GmbH) with ganglion cell side towards the electrodes. The MEA chamber was perfused (flow rate 3 mL/min) with Ames medium, which was oxygenated with a medical-grade gas mixture of 95% oxygen and 5% carbon dioxide at room temperature (22°C). The retinas were left to stabilize from surgical trauma on the MEA perfused with oxygenated Ames medium for at least 30 minutes before being stimulated visually or chemically. A slight suction was applied at the bottom of the perforated MEA to prevent the retina from moving due to the perfusion and to maintain a firm contact with the electrodes throughout the entire duration of the experiment. All sample preparation work and stimulation experiments in this study were conducted under dim red illumination to preserve the light sensitivity of photoreceptors during data collection.

### Experimental Setup

A special experimental platform (schematically illustrated in Supplementary Fig. S4) was built to enable chemical injections into the retina tissue from the photoreceptor (top) side while simultaneously recording the responses of retinal neurons at multiple sites around the injection site on the other (bottom) side contacting the perforated MEA electrodes. The perforated MEA electrodes, each 30 μm in diameter, were laid out in a grid (8×8) pattern with an inter-electrode spacing of 200 μm. The extracellular voltages of activated retinal neurons picked up by 60 electrodes of the perforated MEA were amplified and acquired into a computer through an MEA system with MC-Rack software (MEA1060, Multichannel Systems, GmbH).

Pre-pulled glass micropipettes (10 μm tip diameter and 1 mm outer diameter; World Precision Instruments, Sarasota, FL) held in a standard microelectrode/micropipette holder (QSW-A10P, Warner Instruments, Hamden, CT) were used to both store glutamate chemical and serve as a reference electrode. The pressure port of the micropipette holder was coupled with a pressure injector system (PM-8, Harvard Apparatus, Holliston, MA), which was used to inject pulses of glutamate into the retina tissue. The micropipette tip was filled with glutamate buffered in Ames Medium (including NaCl and KCl for impedance measurements) and electrically coupled to a patch clamp amplifier (Axopatch 200b, Molecular Devices, Sunnyvale, CA) with a silver/silver chloride wire to form an electrode. Deviations in the electrode impedance from the initial impedance, monitored through the patch-clamp amplifier output signal, were used to determine contact of the micropipette tip with the retinal surface as the tip was inserted into the retina and for any blockage of the pipette tip (by extracellular material) during an
experiment. The spatial position of the micropipette tip with reference to fixed reference points on the MEA was controlled by a three-axis, motorized, precision manipulator (MP-285, Sutter Instruments, Novato, CA) with sub-micron positioning accuracy.

The set-up also included a small liquid crystal display (Ruby SVGA Color Microdisplay, 800μ600 pixels, Kopin, Westborough, MA) driven by a computer running Psychophysics Toolbox (PTB-3) in MATLAB (MathWorks Inc., Natick, MA) to enable GWN checkerboard stimulation of the retina from the bottom side and a green (570 nm) light-emitting diode (LED) to stimulate the retina with full-field flashes of light from the top.

To achieve repeatable, high accuracy spatiotemporal modulation and synchronization of the chemical and visual stimulus events and precise positioning of the micropipette tip in the subsurface of the retina in each experiment, all of the main instruments (pressure injector, light sources, and the manipulator) in the set-up were computer controlled via a digital-to-analog DAQ board (PCI-6251, 16-bit, National Instruments) and custom scripts coded in LabView (National Instruments, Austin, TX). In addition, the set-up was also equipped with a stereomicroscope (SMZ 745T, Nikon) supported on a gantry and a digital camera (Moticam 5, Motic, Richmond, BC, Canada) to enable observations of the region near the micropipette during the experiments.

### Stimulation Protocol

Before starting the stimulation protocol, the health of the retina sample following the surgical procedure was monitored by observing the spontaneous and light-evoked responses of RGCs for about 30 minutes. Once stabilized, the retina was investigated according to the following major protocol steps: 1. The locations and extent of RGC RF centers as well as RGC subtypes were identified using responses evoked by a GWN checkerboard stimulus projected onto the RGCs from below, 2. The functional health of the retina was assessed, and in some samples LFP responses were recorded, using responses evoked by a full-field visual stimulus aimed at the photoreceptor-surface from above, and, 3. gSRs were recorded by injecting boluses of glutamate targeted at the RF centers of RGCs, which had robust visual responses, predominantly cells identified as OFF or ON RGCs. The details of the visual and chemical stimuli and the stimulations performed in each of the protocol steps are described below.

The GWN checkerboard stimulus was a grayscale 24×32 pixel image generated with the LCD and focused onto the retina using a camera lens such that the image projected by each checker on the retina measured 100 μm × 100 μm corresponding to a viewing angle of 0.33° × 0.33°. The LCD frame rate was 60 Hz and checkerboard patterns were presented at a rate of 15 Hz. The mean illuminance of the LCD on the epiretinal surface was 4.9 lm/m^2^. An initial 25 min period of GWN checkerboard stimulation was used to quickly characterize and locate RGC RFs for targeted stimulation, followed by a comprehensive 60 min stimulation to better characterize RF properties^37^. The full-field flash stimulus was generated by the green LED and comprised a series of flashes (5 lm/^2^, 1-2 sec ON and 1-2 sec OFF).

L-glutamic acid was mixed with Ames Medium (1 mM; Sigma-Aldrich, St. Louis, MO) and loaded into glass micropipettes to permit glutamate stimulation. The micropipette tip was positioned near target RFs with a micromanipulator (see Supplementary Fig. S5a). Once positioned above a target location, the pipette was lowered until contact with the photoreceptor-surface was detected^16^ through an increase in the impedance of the electrode and then inserted 70 μm below the surface to place the aperture near the OPL. After positioning the pipette, glutamate was injected at the target location using pressure pulses of 0.69 kPa applied through the pressure injector system. Glutamate injection trials at each target location comprised three sets of 30 pulses at a constant interpulse duration of 3 s, where the pulse width within each set was maintained to be 10 ms, 20 ms, and 30 ms, respectively (Supplementary Fig. S5b). Similar glutamate injection trials were repeated at multiple locations on each of the retina samples tested. Periods of spontaneous activity and further full-field flashes were recorded between injections to track the health of retinal neurons during and after glutamate stimulation.

In a subset of recordings (2 of 18 retinas), the frequency of glutamate injections was varied while maintaining a constant pulse duration of 10 ms to investigate the temporal resolution of glutamate stimulation. Interpulse durations ranging between 150 and 4000 ms, resulting in stimulation frequencies ranging from 0.25-6 Hz, were tested. In these stimulation trials, the number of injection cycles was varied in proportion to the stimulation frequency, i.e., fewer (minimum 30) cycles for low frequencies and larger (maximum 200) cycles for high frequencies, in order to collect a sufficiently large quantity of data for comparisons across all stimulation frequencies.

### Data Acquisition

RGC spikes were recorded using MC_Rack (Multichannel Systems, 10 kHz sampling, after high pass filtering, 200 Hz cutoff) with an amplitude threshold of approximately – 16 μV. In a subset of our recordings (9 of 18 retinas), raw voltage data (sampled at 10 kHz with no filtering) were recorded for each electrode in addition to the filtered spike data acquired through MCRack. The raw voltage data were processed in MATLAB by low pass filtering (20 Hz cutoff) and averaged across trials to produce average LFP waveforms^38,39^.

## Data Analysis

### Spike Sorting

Spikes were sorted into individual units using Offline Sorter (Plexon Inc., Dallas, TX) and MATLAB. Offline Sorter was employed to quickly identify units through principal component analysis (PCA) to enable targeted injections at RF centers in subsequent recordings using data from the initial short GWN stimulus. After concluding an experiment, data from all recordings were combined in MATLAB and sorted into units using the wavelet clustering package, Wave_clus, developed by Quiroga et al.^40^. Once sorted, PSTHs were calculated from unit spike trains using Gaussian kernel density estimation to average spike rates across trials^41,42^ with custom MATLAB code. Spike rate amplitudes were calculated as the difference between extrema and the mean spike rate, with negative and positive amplitudes characterizing inhibitory and excitatory phases, respectively (Supplementary Fig. S6a). If responses contained more than one phase with amplitudes exceeding a spike rate threshold of 3 Hz, the largest amplitude was assumed to be the dominant response.

Responsive units were identified by comparing fano factor derived response variables (FRVs) from glutamate-evoked and spontaneous PSTHs as expressed in Eq. (1). Units were deemed responsive to a glutamate stimulus if their glutamate FRV was greater than three standard deviations of its spontaneous FRVs. Responsive units were also required to have spike rate amplitudes of 3 Hz or greater.

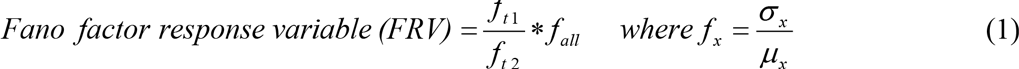

Each fano factor (*f_x_*) is calculated using different parts of the PSTH. The numerator (*f_t1_*) is calculated using the standard deviation (σ) and mean (μ) from the initial segments of the PSTH, usually *t* < 0.5 (0.1 for high frequency stimulation) seconds. The denominator *ft2)* is derived from the remainder of the PSTH (*t* > 0.5 or 0.1 seconds) while the last term (*f_all_*) is the fano factor using all time points. Responsive units are identified by finding glutamate-evoked FRVs that exceed three standard deviations of the cell’s spontaneous FRVs. Implementation of this metric allowed fast detection of transiently responsive units and was insensitive to the basal spiking deviations of individual units. Equation (1) was also used to detect transient LFP responses using the LFP data in place of spike rate data with a threshold of 6 standard deviations (see below).

### LFP Analysis

The raw recorded LFP data, which represent the responses of non-spiking retinal neurons, obtained for full-field flash, glutamate injections, and spontaneous conditions were inverted compared to transretinal ERG waveforms (since the reference electrode was placed above the distal retina). As such, all LFP records shown have been inverted to facilitate comparison between each other and previously reported ERGs^26,43^. A modified FRV (Eq. (1) used in conjunction with LFP data instead of spike rate data) was used to determine responsiveness, where responsive cells displayed glutamate LFP FRVs outside of 6 standard deviations of their spontaneous LFP FRVs.

### Receptive Field Analysis

Following spike sorting, the location and extent of each RGC RF center was characterized using spike-triggered averaging (STA) with the spikes recorded during GWN stimulation utilizing techniques described by Cantrell *et al*.^37^. The 30 GWN frames (2 s) preceding each spike were gathered into a large array to assemble a spike-triggered ensemble (STE). The pixel values of the STE were normalized to a grayscale with limits of -1 (black) to 1 (white). The STA was calculated by performing a vector average of the STE, producing a set of 30 frames representing the average visual stimulus that generated spikes. RGCs with substantial visually-evoked responses yielded STA frames showing large deviations (close to 1 or –1) from a mean background of 0. The frame displaying the largest deviation from background was identified and fitted with a 2D Gaussian to produce an estimated size and location for the RF center. The extent of the RF center was assumed to be the area within a 1-standard deviation ellipse centered on the RF midpoint and a threshold of 6 standard deviations from background was used to eliminate RGCs with weak visually-evoked responses.

### STC-NC Analysis and RGC Subtype Identification

Non-centered spike triggered covariance (STC-NC) analysis was completed following the techniques described by Cantrell *et al*.^31^. STC-NC analysis was initiated by implementing a spatial window of 5×5 pixels on the STE, aligned with the central checker derived from the STA. The windowed STE was then examined with a non-centered PCA to find the eigenvectors with the greatest eigenvalues that maximize the second moment of the STE around zero, thus estimating the linear filter that best predicts the responses elicited. The output of this linear filter was used as an input for a static nonlinearity representing the preference for an RGC to respond nonlinearly to certain stimuli (i.e. ON, OFF, or both) and this combination (a linear-nonlinear model depicted in Supplementary Fig. S6b) was used to predict spike probability. STC-NC analysis identifies RGC subtypes by estimating the static nonlinearity using the measured linear filter (STC-NC eigenvectors) and the observed spike probability. The static nonlinearity was thus calculated as the element-by-element quotient of the output of the linear filter for the STE divided by the linear filter output for all stimulus frames. The resulting static nonlinearity vector represents the probability for RGC spikes in response to light (stimulus standard deviations > 0) and dark (stimulus standard deviations < 0) stimuli presented to the RF. The positive and negative segments of this static nonlinearity vector were integrated to obtain scalar probabilities of responses to light (*P_ON_*) and dark (*P_OFF_*) stimuli. These probabilities were used in Eq. (2) to determine the cell’s bias, which allowed identification of cells as either ON-center (bias ≥ 0.3), OFF-center (bias ≤ -0.3), or ON-OFF-center (–0.3 < bias < 0.3). RGCs with weak light responses and those lacking RFs were unclassified.
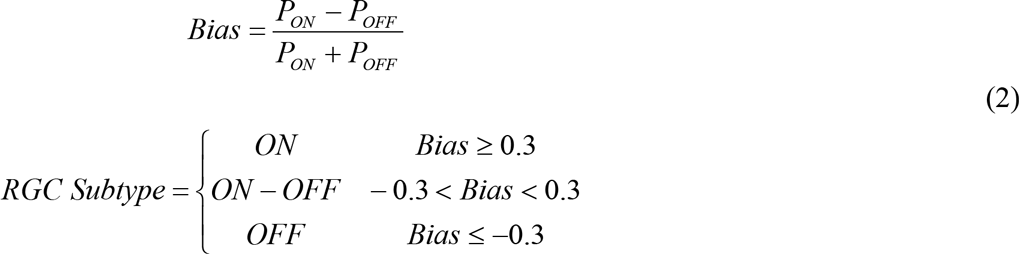

The bias was determined using the probabilities of light (*P_ON_*) or dark (*P_OFF_*) stimuli causing spike esponses, which were calculated from the static nonlinearity. RGCs with positive biases respond strongly to light stimuli in their RF centers while those with negative biases respond strongly to dark. RGCs with biases close to zero respond to both light and dark stimuli.

### Spatial Properties of Glutamate Stimulation

Although glutamate injections were specifically targeted at the RF centers of cells with robust ON- and OFF-center biases, it is very likely that convective diffusion of glutamate resulted in stimulation of surrounding cells as well. Therefore, to estimate the spatial localization of glutamate stimulation, the vectors separating glutamate responsive RGC RF centers from the injection site were measured. These vectors were spatially binned to form a 2D histogram showing the approximate location of responsive RGC RF centers with respect to the injection site. These data were also fit with a 2D Gaussian spread function to produce a more continuous representation of spatial localization.

The spatial properties were further investigated by binning cells based upon the distances separating their RF centers from the injection site using 100 μm wide bins. For each bin, the normalized spike rate amplitude (glutamate-evoked amplitude divided by peak full field flash amplitude) was calculated to enable comparison across cells with widely different basal spiking rates.

## Statistical Analyses

A one-sided, two-proportion *z*-test (α = 0.05) was used to statistically compare the proportions of inhibitory OFF RGC responses to glutamate injected inside the RF centers of cells with the proportion of inhibitory responses to injections outside RF centers (Eq. (3)). A one-sided test was chosen to test the hypothesis that inhibitory OFF responses were predominantly confined to injections inside the RF center. This comparison showed that OFF RGCs with both excitatory and inhibitory responses possessed significantly (*p*-value = 0.026) more inhibitory responses to injections inside their RF centers compared to injections outside (difference in proportions = 0.18; *z*-statistic of 1.94).

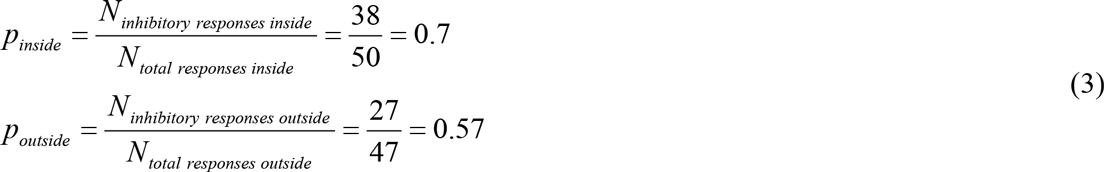

The normalized spike rate amplitudes described in the previous section were analyzed (α = 0.05 ,*p* < 0.0001, Kruskal-Wallis test) to examine the differences between bins. This test was utilized because the normalized spike rate amplitudes were non-normally distributed. The sample sizes of each bin is reported in Supplementary Fig. S2a. A similar test was used to compare the non-normally distributed LFP amplitudes (α = 0.05, *p* = 0.0489, Kruskal-Wallis test) in Supplementary Fig. 2c.

## Acknowledgments

The work presented in the paper was supported by the National Science Foundation, Emerging Frontiers in Research and Innovation (NSF-EFRI) program grant number 0938072. The contents of this paper are solely the responsibility of the authors and do not necessarily represent the official views of the NSF.

### Author Contributions

All authors contributed to the design of the experiments and data analysis. C.M.R. and S.I. developed the experimental setup with oversight from L.S. C.M.R. conducted the experiments, recorded and analyzed the electrophysiological data. J.B.T. oversaw the visual receptive field and electrophysiological data analysis. L.S. initiated and managed the project. All authors discussed the results and prepared the manuscript.

### Additional Information

**Competing financial interests**: The authors declare no competing financial interests.

